# Cell wall remodeling and vesicle trafficking mediate the root clock in Arabidopsis

**DOI:** 10.1101/2020.03.10.985747

**Authors:** Guy Wachsman, Jingyuan Zhang, Miguel A. Moreno-Risueno, Charles T. Anderson, Philip N. Benfey

## Abstract

In Arabidopsis, lateral roots initiate along the primary root in a process preceded by periodic gene expression, a phenomenon known as the root clock. Many genes involved in lateral root initiation have been identified. However, very little is known about the structural changes underlying the initiation process nor about how root clock function is regulated. In genetic screens, we identified the vesicle trafficking regulators, GNOM and its suppressor, AGD3, as critical to root clock function. We show that GNOM is required for the proper distribution of pectin, a mediator of intercellular adhesion, and that pectin esterification state is essential for a functional root clock. We found that in sites of lateral root primordia emergence, both esterified and de-esterified pectin are differentially distributed. Using a reverse genetic approach, we identified significant enrichment of GO terms associated with pectin modifying enzymes in the oscillation zone were the root clock is established. In agreement with a recent study on the function of pectin in pavement cell morphogenesis, our results indicate that the balance between esterified and de-esterified pectin is essential for proper root clock function and the subsequent initiation of lateral root primordia.

## Introduction

Both plants and animals use biological clocks to control developmental processes. In vertebrates, the formation of presumptive vertebrae, known as somitogenesis, is one of the best characterized mechanisms involving periodic gene expression. In plants, prior to the initiation of lateral root primordia (LRP), the auxin response marker, DR5, exhibits periodic expression in a region known as the oscillation zone (OZ) where many other genes are also expressed (*1, 2*). This rhythmic gene expression, known as the root clock, produces pre-branch sites that develop into LRP (*1*). The root clock represents a flexible oscillator, which contributes to the formation of pre-branch sites approximately every six hours. Several genes have been implicated in the initiation of LRP, most of which are related to auxin signaling (e.g., (*3, 4*)) or metabolism (*5*). However, the processes by which these signals are translated into morphological changes including modifications of cell wall structures and tissue adhesion are poorly understood. Moreover, it is unclear which genes contribute to the root clock and whether genes that are involved in structural modifications that allow subsequent lateral root (LR) initiation have a role in establishing the root clock. Recently, the biophysical aspects of LR initiation and the contribution of surrounding tissues to this process have been explored (*6*). These authors showed that auxin-dependent changes in the cellular volume of the endodermis are necessary for the initiation of LRP in the underlying pericycle cells, suggesting a complex interaction between these adjacent layers prior to LR initiation. One candidate protein that putatively mediates signaling between cells and is also involved in LR initiation is GNOM (*7*). GNOM is a trans-Golgi network-localized (*8*) *ADP RIBOSYLATION FACTOR GUANINE NUCLEOTIDE EXCHANGE FACTOR* (*ARF-GEF*; (*9*)) that regulates vesicle trafficking to the plasma membrane. It had been suggested that the reduced LR phenotype of the weak *gnom* allele, *fwr*, is mediated by one of the PIN-FORMED (PIN) proteins. However, the *fwr* mutant can be partially rescued by auxin treatment and has properly localized PIN1 (*7*). In addition, there are no known PIN mutants with LR phenotypes. These data suggest that the lack of LRs in the *gnom* mutant might depend on other components that are not directly involved in auxin signaling. Vesicle trafficking is necessary for the secretion of cell wall components including pectin, hemicelluloses and cell wall modifying enzymes (*10*). Here we show that disrupting GNOM alters the distribution of pectin and blocks the functioning of the root clock. Furthermore, higher levels of esterified and de-esterified pectin are differentially localized around emerging LRP, while alteration of either form of pectin leads to complete loss of pre-branch site formation. Our finding together with a recent paper describing the role of pectin nanofilments in pavement cells development (*11*), suggest that balanced pectin esterification is necessary for proper morphogenesis. Based on these data, we conclude that GNOM-dependent vesicle trafficking mediates the distribution of pectin in the cell wall, which in turn is essential for root clock function and subsequent LR formation.

## Results

The root clock was initially identified through oscillation of DR5 expression in a region known as the OZ near the elongation-differentiation border of the primary root of Arabidopsis (*1, 2*). After the peak expression of DR5-driven luciferase (DR5::LUC) its region of expression shrinks to form a pre-branch site, where subsequently, LRP emerge. Formation of pre-branch sites and LRP initiation are tightly linked: all LRP emerge from pre-branch sites and almost all pre-branch sites develop into LRP (*1*). To better understand the relationships between LR initiation and pre-branch site formation, we examined roots expressing both the DR5::LUC reporter, which marks pre-branch sites, and pCLE44::GFP, which marks early LRP (Fig. S1, A and B; see below for details on the CLE44 marker). In the majority of cases (21/26), the latest DR5::LUC-marked pre-branch site was associated with an early stage LRP. We conclude that LRP initiation starts together or soon after the formation of a pre-branch site, which follows the peak expression of DR5::LUC in the OZ.

To better understand the oscillation mechanism of the root clock, we used RNA-seq to identify genes with elevated expression in the OZ and/or in the most recently formed pre-branch site as compared to their adjacent regions. In a spatial experiment, we collected transverse sections from the OZ, the first pre-branch site, and their flanking regions (Fig. 1A) and performed RNA-seq on these samples. In a complementary, temporal experiment we collected DR5::LUC expressing sections before, during and after peak expression and used them for RNA-seq analysis (Fig. 1B and Supplementary materials). To validate the quality of the data, we compared RNA-seq reads of the luciferase transgene (driven by the DR5 promoter) to luciferase imaging and found a high correlation (Fig. 1, C and D). Multidimensional scaling analysis and other statistical analyses demonstrated that spatial and temporal sections generally form separated clades based on their region or time of collection, respectively (Fig. 1E and S2). To identify putative root clock regulators, we selected genes that were over-represented in the OZ-spatial, OZ-spatial and OZ-temporal, or pre-branch site (pre-branch site in the spatial experiment; Fig. 1, F to I and Table S1, A to F) and used these three lists to perform GO analysis (Table S2, A to C). Some of the most common terms were related to cell wall biogenesis and vesicle trafficking (Fig. 2A). Among these categories, we identified one term for pectin-metabolic process, including genes that code for pectin lyases, pectate lyases, pectin methylesterases (PMEs), and their inhibitors (PMEIs) (p-value=0.005; Fig. 2, B and C and Table S2, A to C). PMEs de-methyl-esterify galacturonic acid residues in the most abundant form of pectin, homogalacturonan (HG). Pectin is synthesized in the Golgi in a highly methyl-esterified form and delivered to the apoplast, where it can be de-methylesterified. Block-wise de-esterified HG chains can be crossed-linked by calcium, leading to stiffening of the cell wall and/or promoting adhesion between cells (*12, 13*), whereas randomly de-esterified HG can be degraded by pectinases, resulting in wall loosening and/or cell separation (*14*–*16*). All eight over-represented OZ PMEs belong to group 1 (*17*) within the type-I class of PMEs (*18*), which contain a PRO domain at the N-terminus that requires preprocessing in the Golgi before secretion to the apoplast (*19*). Another group of genes over-represented in the OZ-spatial dataset is involved in auxin transport and signal transduction. Mutations in five of these genes (LAX3, AUX1, ARF19, IAA14 and IAA28) directly affect LR formation (*3, 20*–*22*). A third group over-represented in the OZ-spatial dataset is involved in the transport and metabolism of, and response to nitrogen-related compounds, (Fig. S3A), all processes that have been shown to affect LR density and outgrowth (*23, 24*). For example, the *TONOPLAST INTEGRAL PROTEIN2;1* (*TIP2;1*), a tonoplast aquaporin, is involved in NH_3_ transport to the vacuole and response to NH_3_ accumulation (*25*). Moreover, TIP2;1 is redundantly required, together with two other TIP proteins, for LR initiation (*26*). We also identified two additional genes that were not included in the dataset since they did not fulfill all logFC=0.5 criteria but still had high expression in the OZ: AMT2;1, an ammonium transporter required for ammonium-dependent LR branching (*24*) and TGA4, a basic leucine zipper transcription factor controlling nitrate-dependent LR initiation (*23*). Finally, our approach also identified 10 genes in a GO term for LR morphogenesis (Fig. S3B), which include auxin transport (LAX3 and AUX1), auxin signaling (ARF11/ARF19, IAA14 and IAA28), exocytosis (EXO70A1) and the promotion of protein degradation by ubiquitin ligase activity (XBAT32). Together, these results suggest that the OZ functions as a transcriptional hub for multiple pathways required for emergence of LRP. To further explore this hypothesis and because structural changes in the walls of LRP cells and the cells of surrounding tissues are likely to occur prior to and during LR initiation, we focused on the group of genes regulating pectin metabolism.

**Fig. 1.**
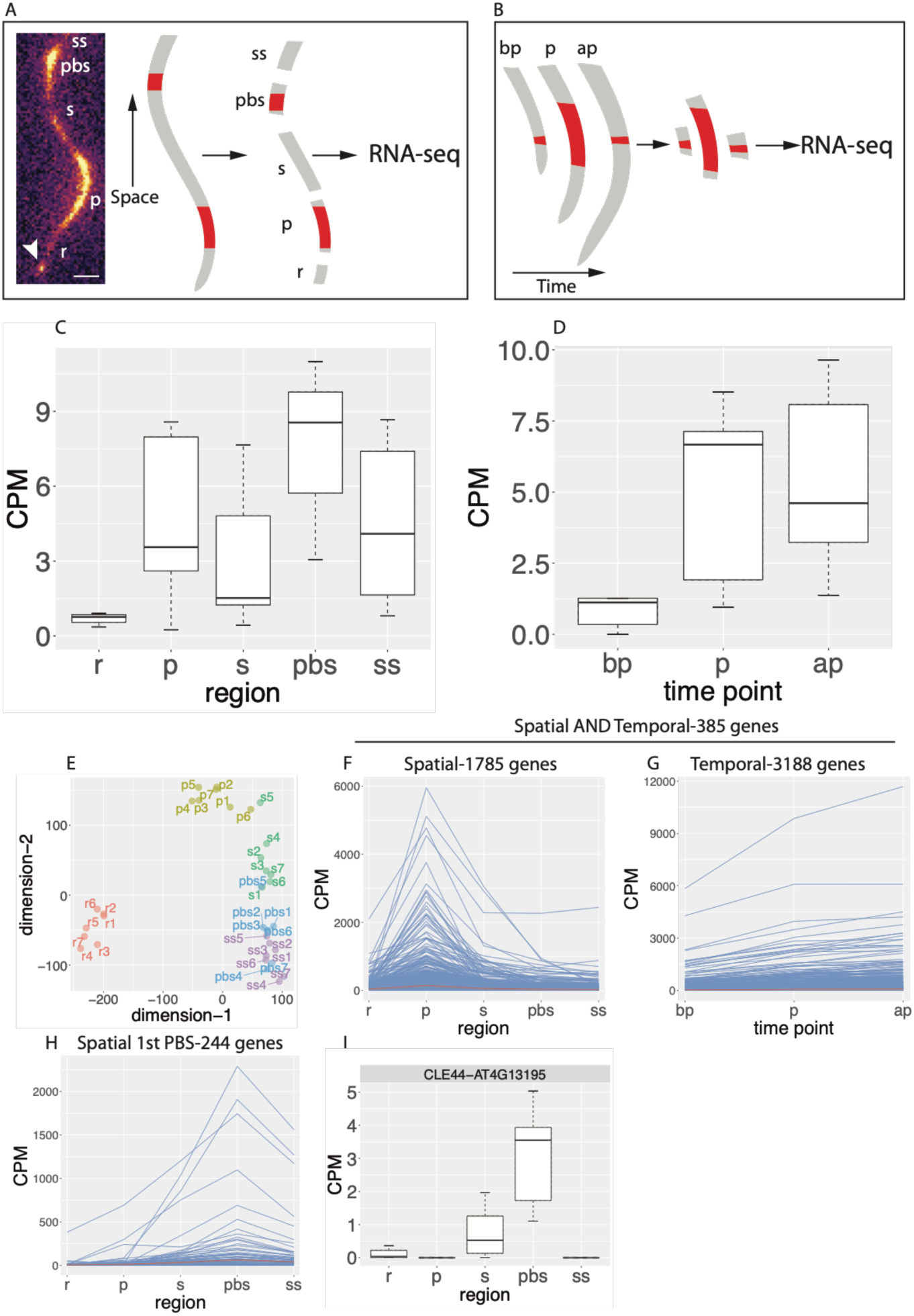
Experimental design and analysis of RNA-seq experiments. **(A)** Spatial experiment. Seven DR5::LUC roots were imaged using the Lumazone system (left). Once a peak expression was detected in the OZ, the root was dissected into five sections: r – a section proximal to the root meristem and below the OZ, p – OZ, s – a section proximal to the OZ and distal to the youngest pre-branch site, pbs – youngest pre-branch site and ss – a section proximal to the pre-branch site. Each section was used to generate a transcriptome profile (35 total). Arrowhead, root tip; scale bar=0.3 mm. **(B)** Same as in (A) but a root section was collected from the OZ either before peak expression (bp), during peak expression (p) or after peak expression (ap). **(C** and **D)** Luciferase CPM RNA reads in the spatial experiment (C) and temporal experiment (D). **(E)** multidimensional scaling plot of the spatial experiment. Note that the regions (r, p, s, pbs and ss) mostly form distinctive clades. **(F)** Expression pattern of all 1785 genes in the spatial experiment with a log_2_ *FC* ≥ 0.5 (*FC* ≥ 1.414) in the p region compared to each of the other zones. **(G)** Expression pattern of all 3188 genes in the temporal experiment with a log_2_ *FC* ≥ 0.5 (*FC* ≥ 1.414) in the p time point (peak Luciferase expression) compared to bp (before peak Luciferase expression). **(H)** Expression pattern of all 244 genes in the spatial experiment with a log_2_ *FC* ≥ 0.5 (*FC* ≥ 1.414) in the pbs zone compared to each of the other zones. **(I)** Expression profile of CLE44 that was later used as a pre-branch site marker. CPM, counts per million.

**Fig. 2.**
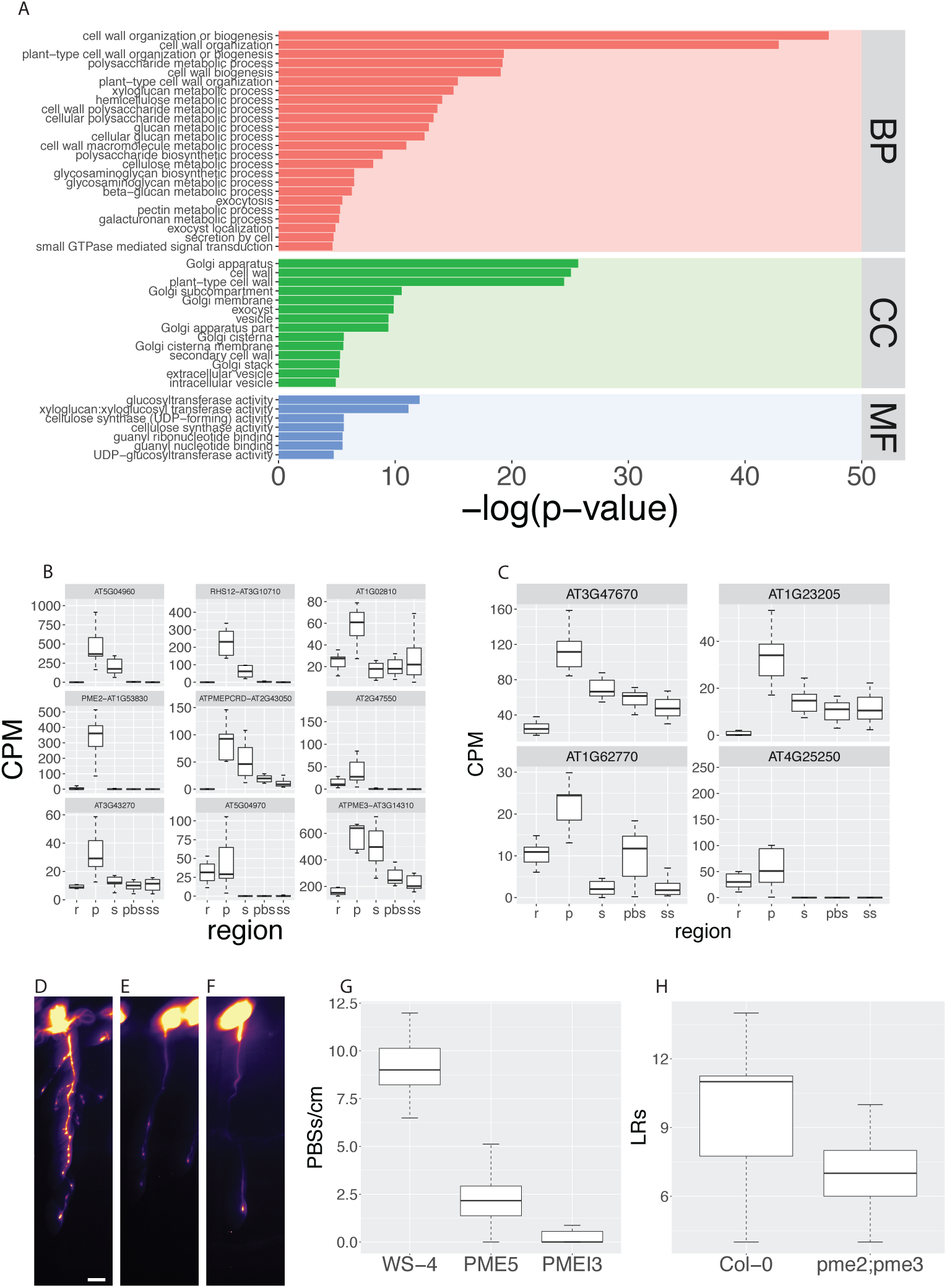
Cell wall, vesicle trafficking and membrane related genes are over-represented in the OZ and pre-branch site and required for its formation. **(A)** GO term analysis of the 1785 genes described in Fig. 1F (BP, biological process; CC, cellular component; MF, molecular function). The plot depicts vesicle trafficking, Golgi, cell wall, small GTPsase mediated signal transduction and carbohydrate metabolism related GO terms with p-value≤0.01 (hypergeometric test using the gProfiler R package). **(B** and **C)** Expression pattern of nine PME (B) and four PMEI (C) genes in the spatial experiment. **(D** to **F)** pCLE44::LUC expression in WT (WS-4, (D)), PME5 overexpressing (E) and PMEI3 overexpressing (F) seedlings under EtOH inducible conditions. Scalebar=0.2 cm. **(G)** Quantification of pre-branch site number (D to F). p-value<10^−14^ (Wilcoxon rank sum test). **(H)** LR number of *pme2*;*pme3* compared to Col-0 WT. p-value=0.007 (Wilcoxon rank sum test).

PME5 and a PMEI3 have been implicated in phyllotaxis (*27*). Shoot apical meristems of PME5 over-expressing plants produce ectopic lateral shoot primordia, whereas induction of PMEI3 leads to a PIN-like apical meristem and suppression of lateral organs. To determine whether changing the esterification state of pectin affects rhyzotaxis as well, we overexpressed PME5 and PMEI3 and in both cases found a severe reduction in the number of PBSs as shown by the pCLE44::LUC and DR5::LUC markers (Fig. 2, D to G and Fig. S4, A to D) as well as a decrease in LR number (Fig. S4, E to G). This result is in agreement with a recent paper showing that overexpression of either of the genes leads to decreased width variability in the anticlinal wall of pavement cells and reduced lobe formation (*11*). Based on our RNA-seq results, we identified two PME genes, whose mutants *pme2* and *pme3* as well as the double mutant combination showed reduced numbers of LRs (Fig. 2H and Fig. S4, H and I). Since PME processivity determines the degree of cell wall stiffness (*18*), which might impinge on both wall expansibility and cell-to-cell adhesion, the effects of ectopic PME and PMEI expression together with the specific expression and functional requirement of the pectin modifying enzymes suggest that the balance between pectin esterification and de-esterification underpins the functionality of the root clock.

To complement our reverse genetics approach, we performed a genetic screen of mutagenized DR5::LUC seeds looking for mutants with an altered root clock. We identified six mutations with modified root clock function (Table S3A) and mapped the affected genes using the SIMPLE pipeline (*28*). Two independent mutations were found in *ROOT HAIR DEFECTIVE 3 (RHD3*), one each in *GNOM, SHORTROOT, GLUTHATIONE REDUCTASE 2* and *SECA1* (Fig. 3A and Table S3). Since three of the six mutations were in genes that code for vesicle trafficking-related proteins (RHD3 and GNOM), this suggested that vesicle trafficking might be important for establishing and/or maintaining root clock activity. Seedlings of the weak *gnom*^*184*^ allele that we identified are slightly smaller than WT with neither LRs nor pre-branch sites and have mild root hair and anthocyanin accumulation phenotypes (Fig. 3 and Fig. S5, A to I). Since the *gnom*^*184*^ mutation is a missense arginine to glycine substitution in the Sec7 domain, which mainly affects the root clock and LR development, it is likely that this process is the most sensitive to GNOM-dependent vesicle trafficking. To better understand the phenotype of *gnom*^*184*^, we grew WT seedlings on Brefeldin A (BFA), a vesicle trafficking inhibitor that targets a subset of plant ARF-GEFs, including GNOM (*29*). Seedlings grown on low concentrations (3-5μM) of BFA have a similar phenotype to *gnom*^*184*^ with no LRs (Fig. S5I and (*29*)). A single amino acid substitution in the catalytic Sec7 domain of GNOM can convert WT seedlings to BFA resistance (*29*) resulting in a mutant developing LRs upon BFA treatment, suggesting that BFA directly and specifically targets the GNOM protein to inhibit LR formation. To test whether BFA has an effect on the root clock, we imaged DR5::LUC and pCLE44::LUC (which was identified as a pre-branch site expressed gene in the RNA-seq experiment; Fig. 1I) expressing seedlings after BFA treatment. The expression pattern of these markers was strongly altered, phenocopying the *gnom*^*184*^ allele (Fig 3, B and C and Fig. S5, G and H). Next, we sought to identify other components of the GNOM-dependent root clock pathway through a suppressor screen. Since BFA is a GNOM-specific inhibitor in the LR context, we used the same mutagenized DR5::LUC population and screened for BFA resistant seedlings that could develop LRs under BFA treatment conditions. We identified two BFA insensitive mutants (Table S3) displaying both LRs and pre-branch sites upon BFA treatment (Fig. 3, B and C and Fig. S5, G to M). The first affected gene, *LEUCINE CARBOXYLMETHYLTRANSFERASE1/SUPPRESSOR OF BRI1* (*LCMT*/*SBI1*) is involved in the termination of the brassinosteroid signaling cascade during endocytosis by promoting degradation of the hormone receptor BRI1 (*30*). The second gene, *ARF-GTP ACTIVATING PROTEIN DOMAIN* (*AGD3*), is required for scission during vesicle formation (*31*). GNOM is an ARF-GEF (ARF Guanine nucleotide exchange factor), which facilitates the exchange of GDP to GTP leading to membrane recruitment of the ARF protein. AGD3 acts in the opposite pathway by activating ARF-dependent GTP hydrolysis, which allows vesicle budding and eventually the dissociation of ARF-GDP from the membrane (*32*). When introduced into the *gnom*^*184*^ background, our mutant *agd3* allele (*agd3*^*26*^) partially rescued the loss of root clock function and LR phenotype (Fig. 3, B and C and Fig. S5, G to I). These results indicate that the vesicle trafficking circuit of GNOM and AGD3 is specifically required for the initiation of LRP and the functioning of the root clock. Since pectic HG is synthesized and methyl-esterified in the Golgi and then secreted to the cell wall where it can be de-esterified by PMEs, we sought to determine whether its involvement in the root clock and LRP initiation requires GNOM-dependent vesicle trafficking from the Golgi to the apoplast. To test this, we labeled three HG forms: de-esterified and Ca^2+^ cross-linked (*33*), low-methyl-esterified, and high-methyl-esterified (*34*), in WT and *gnom*^*184*^ using immunohistochemistry on root cross-sections. The mean intensity signal of HG in all three experiments did not differ between WT and *gnom*^*184*^ (Fig. S6), however the cell wall area occupied by the low-esterified form was significantly lower in *gnom*^*184*^ than in WT (p=0.017, Fig. 4A). It was previously shown that a severe *gnom* allele alters the distribution of de-esterified HG in the cell wall (*35*). Since the localization of de-esterified pectin is altered in *gnom*^*184*^ and the boundary between the pericycle and the endodermis is relevant for the emergence of LRP, we focused our comparison of de-esterified HG distribution on pericycle/endodermis junctions. Using an antibody that preferentially binds low-esterified HG (*36*), transmission electron microscopy identified two types of gold particle distribution in three-way pericycle/endodermis junctions (junctions of two pericycle cells and one endodermis cell or one pericycle cell and two endodermis cells). “Closed” junctions had HG occupying the entire three-cell junction space, whereas “open” junctions were particle-free (Fig. 4, B and C). We measured the gold particle density (Supplementary materials) and found that it was significantly higher in *gnom*^*184*^ junctions as compared to WT (p=0.001, Fig. 4D). This indicates that the *gnom*^*184*^ mutant tends to have more “closed” junctions while WT has a higher abundance of “open” junctions. We also performed a more qualitative comparison by directly counting the number of open, closed and ambiguous junctions based on their overall appearance and found that *gnom*^*184*^ had a significantly lower number of open junctions and a higher number of closed junctions as compared to WT (p<0.002, Fig. S7A). Interestingly anticlinal cell walls separating adjacent endodermis cells where the Casparian strip forms, were signal-free in both the immunohistochemistry and immunogold experiments and the cell wall appeared squashed (Fig. 4C and Fig S6, A to C). We conclude that GNOM is required for proper distribution of de-esterified HG in the pericycle/endodermis junctions and that this distribution might be important for pre-branch site formation and the initiation of LRP. If accurate localization of de-esterified pectin is required for LRP initiation, we hypothesized that there would be differential distribution of esterified pectin at sites of emerging LRP. To test this hypothesis, we labeled transverse sections with antibodies against de-esterified and esterified HG (*36*) at sites of LRP initiation. We found that both the pericycle-endodermis and the endodermis-cortex circumference boundaries adjacent to the LRP have significantly lower levels of low-esterified HG as compared to the opposite side (Fig. 4, E and G). In contrast, we identified higher levels of high-esterified HG in the endodermis-cortex boundary (but not in the pericycle-endodermis) adjacent to the LRP as compared to the opposite side (Fig. 4, F and H). These results show that de-esterified HG, which is usually associated with increased stiffness of the cell wall and tighter cell-cell adhesion, is eliminated near emerging LRP and that pectin subtypes are differentially distributed at sites of initiating LRP. Taken together, our results indicate that the process of secreting and modifying pectin is critical to the function of the root clock and subsequent LR formation. Given that LRP emerge by pushing through overlying cell layers, loss of de-esterified pectin could reduce cell adhesion at pre-branch sites, providing a competence to form LRP.

**Fig. 3.**
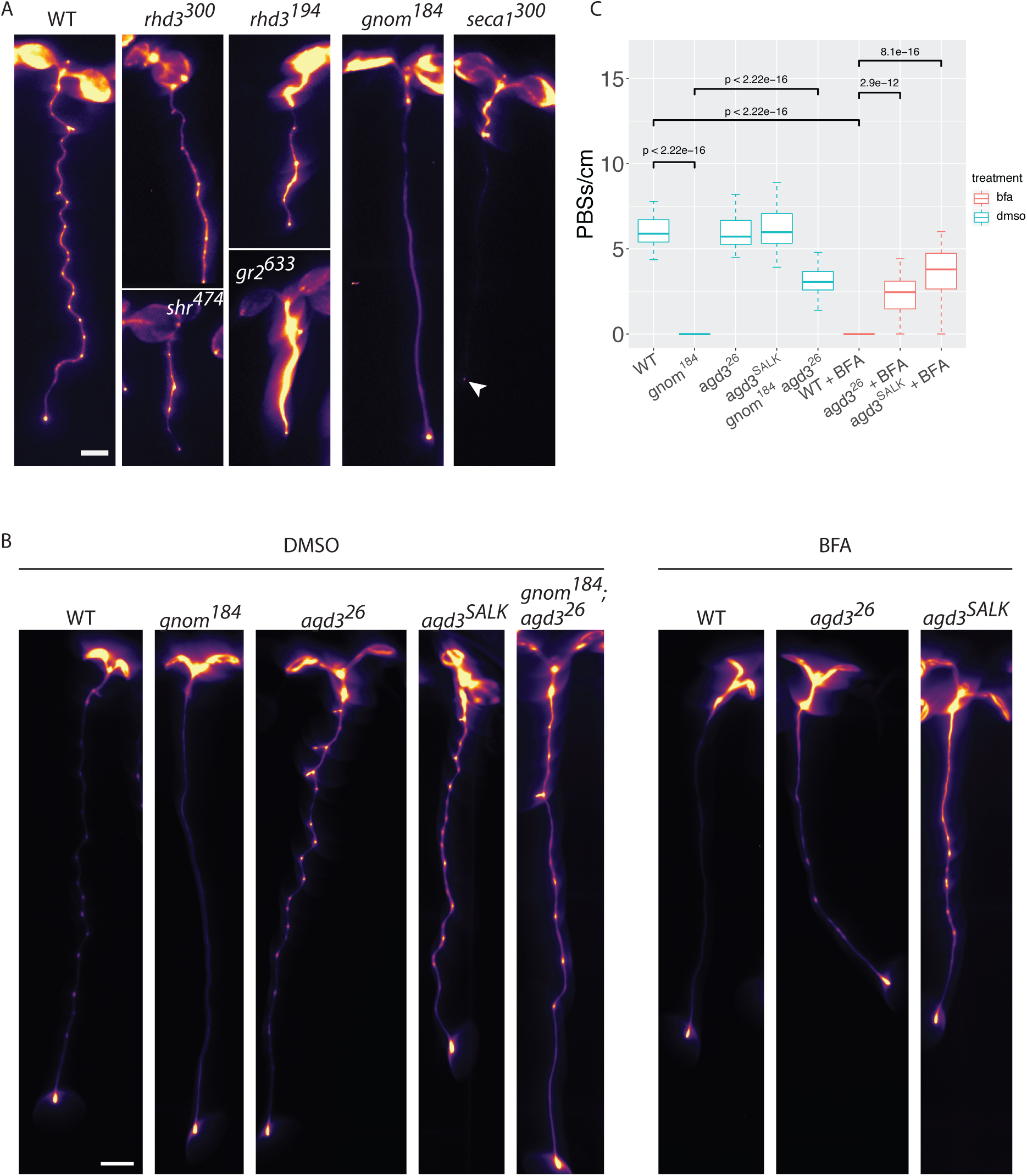
Perturbation of vesicle trafficking affects the root clock. **(A)** Root clock phenotype of EMS mutants. Line name is shown as superscript and gene name in lowercase letters. arrowhead, DR5::LUC expression in the root tip. **(B)** Expression of pCLE44::LUC seedlings in different genotypes under mock (DMSO, left five panels) or BFA treatment (right three panels). **(C)** Quantification of pCLE44::LUC pre-branch site number in *gnom*^*184*^, its suppressor *agd3* (SALK and the EMS mutant *agd3*^*26*^) with or without 3 μM BFA (t-test). Scalebar=0.2 cm.

**Fig. 4.**
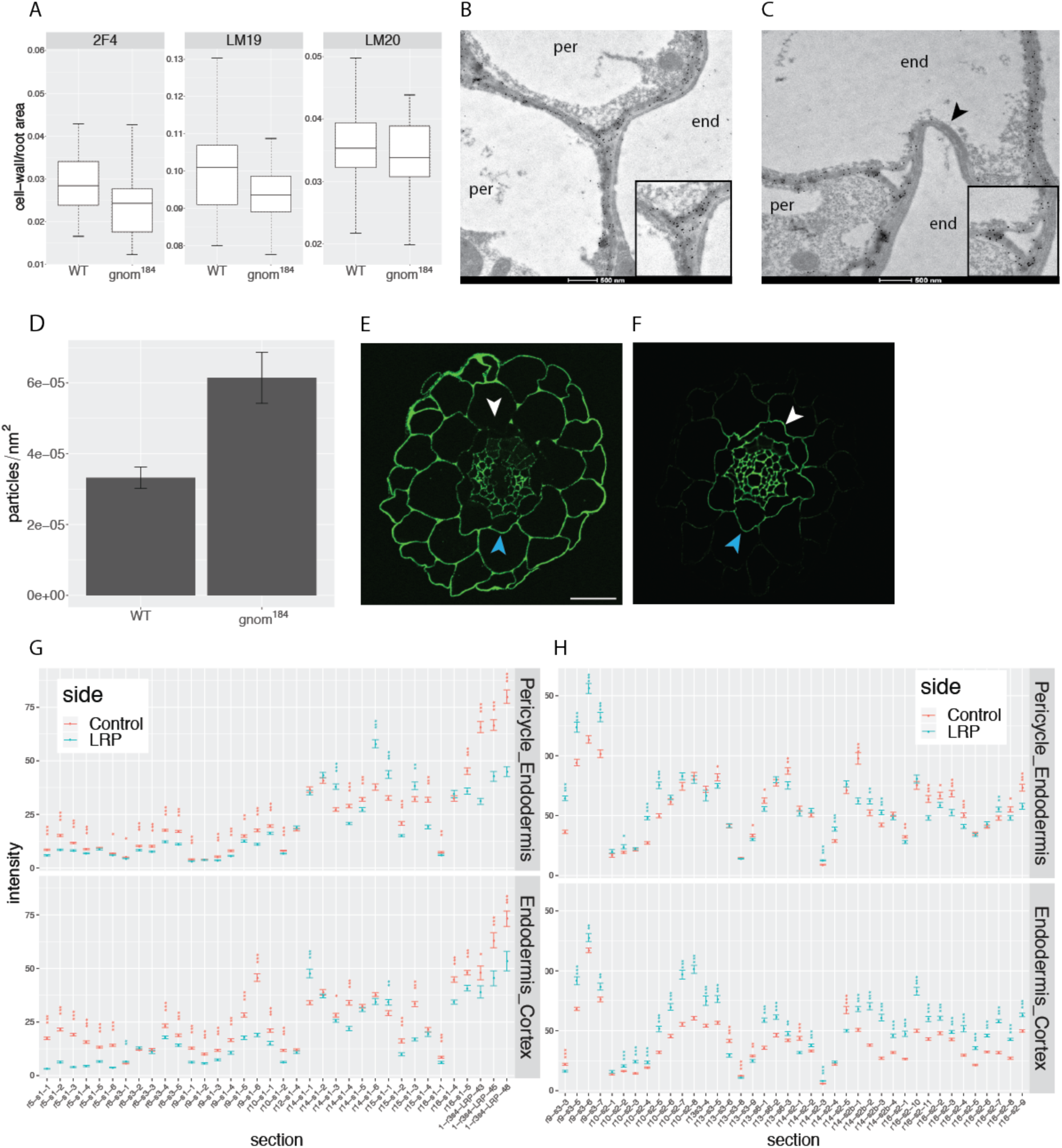
Low-esterified HG distribution is affected in *gnom*^*184*^ and negatively correlates with sites of LR inception. **(A)** Immunohistochemistry quantification of cell wall area per root cross section area for each antibody (Ab). p-value_LM19_=0.017 (t-test). **(B** and **C)** TEM images of LM19 immunogold localization. De-esterified HG is marked by round black gold particles. Cell type (end, endodermis or per, pericycle) is indicated in the cytoplasmic region of each cell. Insets show the three-way “open” (B) and “closed” (C) cell junction types. Note the collapsed and antibody-free endodermis-endodermis cell wall marked by arrowhead. **(D)** Quantification of gold particles in pericycle-endodermis junctions; genotypes show different particle density (p-value=2.2*10^−16^, Wilcoxon Rank Sum test). **(E** and **F)** Immunohistochemistry with LM19 (E) and LM20 (F) of root cross-sections in regions of emerging LRP. Note the enhanced (blue arrowhead) signal in the wall opposite the LRP (white arrowhead) with the de-esterified HG Ab (E) and the opposite trend with the anti-esterified HG Ab (F). Scalebar=20 μm. **(G** and **H)** Quantification of (E) and (F) respectively. For each root section (x-axis), mean pixel intensity was measured in the pericycle-endodermis (upper panels) or endodermis-cortex (lower panels) cell wall border in the LR and opposite side (*, p-value≤0.05; **, p-value≤0.01; ***, p-value≤0.001). Asterisk color matches the side with the higher intensity.

## Discussion

A minimum requirement for an oscillating system is delayed feedback (*37*), which can produce a periodic pattern with as few as two components. However, the somitogenesis clock requires the function of multiple components arranged in three complex networks (e.g., (*38*)). The Notch1 signaling pathway, one of the three pathways governing somitogenesis, requires vesicle recycling through the trans Golgi network (*39*). This indicates that the secretory pathway can play an important role in generating and/or maintaining rhythmic behavior. In Arabidopsis, more than 30 genes have been implicated in LR development, most of which are related to auxin signaling, transport, and homeostasis (*40*). However, very little is known about genes that control pre-branch site formation leading to the positioning of LRs. A recent study showed that endodermis cells overlying early LRP undergo volume reduction prior to emergence (*6*) and that these morphological changes are necessary for the first asymmetric division that marks the LR. Here we show that a functional root clock as well as LRP initiation involves differential distribution of esterified and de-esterified pectin, with de-esterified pectin levels reduced in cells that overlie primordia and increased on the opposite side of the primordia. This distribution could allow the physical emergence of LRP while preventing a similar event to take place on the opposite side of the root. Additional support for this hypothesis that a balanced esterification level is essential for both the oscillation that forms the root clock and initiation of LRP, is shown by the overexpression phenotype of either PME5 or PMEI3 which confers similar phenotypes in the root (this study) and in pavement cells (*11*). The fact that two proteins (GNOM and AGD3) with opposing functionalities also have an opposite phenotype, provides additional evidence for the importance of balanced pectin components. Finally, positing feedback between wall composition and exocytosis was the only way of achieving concordance between experiment and model of the effect of PME and PMEI overexpression on pectin’s role in cotyledon cell morphogenesis. Together, these results suggest that cell wall changes in non-pericycle cells are important for pre-branch site and LR formation as positive regulators of this process. One of the outstanding questions is how the status of extracellular pectin is conveyed to the cell in order to initiate LR formation. A recent study showed that the crRLK kinase, FERONIA, which is required for indentation in pavement cells, directly binds pectin (*41*). In our data, four crRLKs are highly expressed in the OZ (Table S1C) which raises the possibility of pectin-dependent involvement of these receptors in LR initiation. Although pectin modification is necessary for lateral organ formation both in the shoot (*27*) and in the root, pectin modification seems to have different effects in these two poles. In the shoot, methyl-esterification and de-esterification have opposite effects on lateral organ formation, while in the root, our results indicate that a spatiotemporally regulated balance between esterified and de-esterified pectin is essential for lateral organ formation. It is possible that the different developmental contexts and tissue locations in which lateral primordia are formed require different modes of PME activity, with differing developmental consequences. The finding that changes in secretory processes and cell wall components lead to malfunction of the root clock suggests a complex feedback mechanism that relates processes required for LR emergence to the oscillatory process that leads to pre-branch site formation.

## Supporting information

cell_wall_inten_ijf.txt

fiji-count_gold_particles_per-area.txt

threshold_measure_wall_intensity_raul.txt

counts_spatial.txt

counts_temporal.txt

TAIR10_functional_descriptions.txt

Table S1. RNA-seq read count results

Table S2. Results of GO term analysis

Table S3. EMS screens results

Ab_lrp.R

TEM_measurement.R

analysis_spatial.R

analysis_temporal.R

gprofiler.R

luc_count_spatial.sh

pipeline_spatial.sh

pipeline_temporal (including luc reads and binding).sh

supplemental figures and supplemental file description

## Acknowledgements

Thanks to Carmen Wilson and Yue Rui for technical lab support and advice, Alexis Peaucelle for seeds, Michelle Plue (Duke, SMif), Ricardo Vancini and Hal Mekeel (Duke Pathology) and Kathleen Pryer for assistance with immuno-tissue processing and TEM microscopy. Thanks also to the Duke Center for Genomic and Computational Biology and to Isaiah W. Taylor, Rachel Shahan, Joop Vermeer and Joanna K. Polko for critical review of the manuscript.

## Author contributions

GW and PNB conceived the project. GW and JZ performed the experiments. MAM assisted with mapping. CTA provided supervision and resources for immunohistochemistry. GW wrote an initial draft. PNB, JZ, MAM and CTA reviewed and edited the manuscript.

## Data and materials availability

All materials are available upon request. The RNA-seq and DNA-seq datasets are available in the SRA repository with the accession numbers PRJNA478564 and PRJNA490692, respectively.

## Materials and Methods

### Plant material and growth conditions

PMEI3 and PME5 overexpression and DR5::LUC lines were described elsewhere (*1, 27*). The pCLE44::LUC and pCLE44::GFP lines were generated by fusing the 5144 bps promoter region of CLE44 to the Luciferase or GFP coding regions in a three-way Gateway reaction. The *agd3* mutant allele SALK_140826, *pme2* (GK-835A09), *pme3* (GK-002A10) and *pme2;pme3* (*42*) were acquired from the ABRC. All plants were grown on 0.5GM medium containing 1.1 gr Murashige and Skoog basal salts (in 500 ml ddH_2_O), 1% sucrose, 1% plant agar and 5 ml (in 500 ml ddH_2_O) MES (50 gr/l, pH=5.8 with KOH). BFA was prepared as 50 mM in DMSO stock solution and 30 μl were added to 0.5 l 0.5GM media to prepare 3 μM BFA medium; 0.5GM with 30 μl DMSO in 0.5 l medium was used as control. For RNA-seq experiments we used 1.2% agar instead of 1% and supplemented the medium with 0.4 mM Luciferin from a 400 mM (in H_2_O) stock solution. Plants were stratified for 2 d in a 4°C dark room and grown vertically for 4-10 days under long-day light conditions. For overexpression of PME5 or PMEI3, WS-4 plants or plants carrying the inducible construct were plated and grown on 0.5GM plates as described above followed daily application of 25 μl EtOH 3-4 cm from the root tip starting two days post germination.

### RNA and RNA-seq library preparation

Each root region (spatial experiment) or time point (temporal experiment) had seven or five biological repeats, respectively. Each root section was placed in 5 ml H_2_O, froze in liquid nitrogen and ground with Fisherbrand pellet pestle cordless motor. RNA was prepared using the RNAzol reagent according to the manufacturer’s instructions. In short, disrupted tissue was mixed with 0.2 ml RNAzol and 80μl H_2_O for 5-10 mins at room temperature and centrifuged at 12,000 g for 15 mins. Upper supernatant was transferred to a new tube, 1.25μl (0.5% of the supernatant volume) of 4-bromoanisole was added, mixed and stored at room temperature for 3-5 mins, then centrifuged at 12,000 g for 10 mins at 4°C. Supernatant was transferred to a new tube, 1 μl Glycoblue was added and mixed with 300 μl isopropanol. RNA was precipitated overnight and centrifuged for 30 mins at 4°C. The pellet was washed twice with 75% EtOH by releasing it from the tube wall, air-dried, and resuspended in 10 μl H_2_O. RNA-seq libraries were prepared using the Smart-seq2 protocol (*43*).

### Computational and statistical analysis

Illumina HiSeq 2000 reads were aligned and counted using the pipeline_spatial.sh (whole RNA) and luc_count_spatial.sh (Luciferase RNA read count) bash files for the spatial experiment, and the pipeline_temporal (including luc reads and binding).sh file for the temporal experiment. These code files generate read-count for the TAIR10 gene list (TAIR10_functional_descriptions.txt) of all libraries (counts_spatial.txt and counts_temporal.txt). The two read-count files where then used for statistical analysis and plotting using the R code files analysis_spatial.R, analysis_temporal.R and gprofiler.R. To select for genes that are highly expressed in the OZ-spatial, we determined a logFC=0.5 between the p region and each of the other regions (r, s, pbs and ss) using the contrast function of the edgeR package. For the temporal experiment, we required a logFC=0.5 when contrasting the bp and p time points in order for a gene to be considered as having increased expression. Only GO terms with p-value<0.01 for the first two gene lists (Table S1, C and E) and p-value<0.05 for the third list (Table S1F) were considered. All measurements of LR and pre-branch site number were performed with n≥16. For immunohistochemistry measurements, we used our Fiji macro threshold_measure_wall_intensity_raul.txt to select and measure the cell wall area in regions with Abs localization while the cross-section area was calculated manually using the circle selection tool in Fiji. The cell wall area of each section was integrated and divided by the root cross section area to get the ratio (related to Fig. 4A). We used the Fiji macro cell_wall_inten_ijf.txt to select cell wall regions and the raw integrated density. These two values were summed, and their ratio was calculated to get the grey level per pixel (related to Fig. S6). To measure the intensity of LM19 and LM20 in LRP and control side borders (Fig. 4, G and H), we drew a line along the bordering cell wall, and used the list option under the plot profile Fiji analyze menu. We used Ab_lrp.R code file to generate the plots and calculate p-values of each Ab. For each Ab in each genotype n_sections_>23 and n_roots_=5. For counting and measuring the number of pectin-marked gold particles per nm we used our Fiji macro fiji-count_gold_particles_per-area.txt and R code TEM_measurement.R, respectively. For each genotype, n_roots_≥4, n_sections_≥9 and n_junctions_≥105.

### Fixation, dehydration, infiltration and labeling

3-5 days post germination seedlings were fixed for 2.5 h in fixative composed of 1.6% (w/v) paraformaldehyde and 0.2% (w/v) glutaraldehyde in 25 mM sodium phosphate, pH=7.1. We used polypropylene plates (24 well, Thomas Scientific, 1185U58+1185U62) and insert (PELCO Prep-Eze™ 24-wellplate Insert; Ted-Pella) to avoid chemical interaction with the resin. Fixed roots were rinsed with sodium phosphate buffer, pH=7.2 twice for 15 min each, and with water twice for 15 min each. Samples were dehydrated at room temp through a graded ethanol series (20, 35, 50, 62, 75, 85, 95, 100, 100, 100% [v/v], (with the last 100% step including 0.5% [w/v] filtered Fast Green FCF) for 30 min at each step.

Samples were Infiltrated with a graded series of cold LR White embedding resin (Ted Pella) at 4°C (33% and 66% resin in 100% ethanol, 24 h each, followed by three changes of 100% resin, for 24 h each). Samples were cured in gelatin capsules, 100% resin, for 48 h at 60°C. For immunofluorescence, 2 μm sections were prepared using a Reichert-Jung Ultracut E microtome, labeled as previously described (*44*) and visualized using a Zeiss confocal microscope. Immunogold labeling was performed as described (*45, 46*).

