## Supplementary material for "Cell wall remodeling and vesicle trafficking mediate the root clock in Arabidopsis": luc_count_spatial.sh

#!/bin/bash
#281013_root3
num=3 #root number
ROOT_DATE=281013
ROOT=281013_root$num #assign root and the variable "num", e.g., 030314_$num
SECTIONS=([1]="R$num" [2]="P$num" [3]="S$num" [4]="PBS$num" [5]="SS$num") #this is where the fastq folders are
INPUT=/rdata/ngseq/original_data/rna/illumina/Guy/$ROOT #this is the parent directory of the folders "sections"
DATE=$(date +%d%m%y)
OUT_LUC=/rdata/ngseq/Playground/guy/tophat2/$ROOT/$DATE"luc"
LUCINDEX=/rdata/ngseq/Playground/guy/bowtie2/luc_index/luc_index

#creating the directories . Note! if running twice at the same day, the first run will be overiden
for i in ${SECTIONS[*]}; do
	mkdir -p $OUT_LUC/$i
done 

#running tophat
for l in ${SECTIONS[*]}; do
	tophat2 --no-novel-junc -o $OUT_LUC/$l $LUCINDEX $INPUT/$l/fastq/R1.fastq 2>&1 | tee -a $OUT_LUC/$l/$l.log &
done
wait


# rorks w/ one thred: 
#tophat2 --no-novel-junc option -o /rdata/ngseq/Playground/guy/tophat2/030314_root4/130814luc/P4 /rdata/ngseq/Playground/guy/bowtie2/luc_index/luc_index /rdata/ngseq/original_data/rna/illumina/Guy/030314_root4/P4/fastq/R1.fastq 2>&1 | tee -a /rdata/ngseq/Playground/guy/tophat2/030314_root4/130814luc/P4/P4.log
#030314_root2
#030314_root4

#tophat2 --no-novel-junc -o /rdata/ngseq/Playground/guy/tophat2/030314_root4/130814luc/P4 /rdata/ngseq/Playground/guy/bowtie2/luc_index/luc_index /rdata/ngseq/original_data/rna/illumina/Guy/030314_root4/P4/fastq/R1.fastq 2>&1 | tee -a /rdata/ngseq/Playground/guy/tophat2/030314_root4/130814luc/P4/P4.log

#--keep-tmp
