## Supplementary material for "Cell wall remodeling and vesicle trafficking mediate the root clock in Arabidopsis": pipeline_spatial.sh

#!/bin/bash 

#ROOT_DATE=03122018 #need to fill for script - i.d. experiment
DATE=$(date +%d%m%y)
SECTIONS=([1]= [2]= [3]=) #this is the folder names where the fastq files are; define your list of fastq files between the "" and add elements as needed (containing R*.fastq files)
INPUT=/path/to/fastq/files #this is the parent directory of the folders "sections" in each section folder there should be a fastq folder with R1.fastq, R2.fastq or both files
OUTPUT=/path/to/output/folder/$DATE
BT2INDEX=/path/to/bowtie2/index/file_base_name #i.e, the folder "index" should contain the following files: file_base_name.1.bt2, file_base_name.2.bt2, file_base_name.3.bt2, file_base_name.4.bt2, file_base_name.fa, file_base_name.rev.1.bt2, file_base_name.rev.2.bt2
TH2Index=/path/to/Tophat2/index/file_base_name
gtf_file=/path/to/gtf/file
python=pythonX #make sure you have the correct python and necessary packages (https://htseq.readthedocs.io/en/master/install.html); if not, you can locally install it based on this: (http://thelazylog.com/install-python-as-local-user-on-linux/); the pip installation will work w/o errors only after adding "python" as an alias to the .bash file so when pip installation calls "python" it sees the locally installed and not the system python (e.g., 2.6.6)

#creating the directories. Note! if running twice at the same day, the first run will be overiden
for i in ${SECTIONS[@]}; do
	mkdir -p $OUTPUT/$i
done 


#running tophat, only one that looks for splicing files basename is R1 and/or R2 ends with .fastq
for l in ${SECTIONS[@]}; do
	tophat2 -N 1 -o $OUTPUT/$l -i 10 -p 2 --segment-mismatches 3 --min-segment-intron 10 --transcriptome-index=$TH2Index $BT2INDEX $INPUT/$l/R*.fastq 2>&1 | tee -a $OUTPUT/$l/${l}.log &
done
wait


#sorting by read name with samtools 0.1.19; Note, the options and syntax for Samtools 1.x are different!
for m in ${SECTIONS[@]}; do
	samtools sort -n $OUTPUT/$m/accepted_hits.bam $OUTPUT/$m/accepted_hits_sort &
done
wait

#converting to sam files
#for n in ${SECTIONS[@]}; do
	#samtools view $OUTPUT/$n/accepted_hits_sort.bam > $OUTPUT/$n/accepted_hits_sort.sam &
#done
#wait

#counting w/ HTSeq; python and pip can be installed for a specific user using this link (http://thelazylog.com/install-python-as-local-user-on-linux/); before installing pip, create an alias for the local python in the .bash file do the call for python will be directed to the local user. HTSeq in installed (together w/ two other packages) based on this: https://htseq.readthedocs.io/en/master/install.html; for some reason it is invisible when executing help("modules") but it can be imported and used. 
for p in ${SECTIONS[@]}; do
	$python -m HTSeq.scripts.count -f bam -r name -s no -t exon -i gene_id -m intersection-strict $OUTPUT/$p/accepted_hits_sort.bam $gtf_file > $OUTPUT/$p/${p}-1.txt &
done
wait

#Creating edgeR compatible text file (removing the last 5 summerizing records that HTSeq prints)
for q in ${SECTIONS[@]}; do
	awk 'NR<=33602' $OUTPUT/$q/${q}-1.txt > $OUTPUT/$q/${q}.txt & # this command will print all first 33602 (all At#) and redirect it to a new file
done


######################################################################################################
#binding all counts to a single file
LENGTH=$((${#SECTIONS[@]}*2))
COL1=("1," `seq -s, 2 2 $LENGTH`)
#FIELD=$(IFS=, echo "${COL1[@]}")
#ALL_COLS=$(IFS=, echo "1" echo "${COLUMNS[@]}")
COL2=`echo ${COL1[@]} | sed 's/ //g'`
printf "%s\t%s\n" "At_num" "${SECTIONS[*]}" > $OUTPUT/counts.txt

for i in ${SECTIONS[@]}; do echo $OUTPUT/$i/${i}.txt; done > $OUTPUT/dirs.txt
files=`cat $OUTPUT/dirs.txt`
pr -mts $files | cut -f${COL2[@]} >> $OUTPUT/counts.txt
######################################################################################################
