## Supplementary material for "Cell wall remodeling and vesicle trafficking mediate the root clock in Arabidopsis": pipeline_temporal (including luc reads and binding).sh

#!/bin/bash
# to run (from tophat2 directory): /rdata/ngseq/Playground/guy/Notes/Tophat_play_pipeline.sh

#num=3 #root number
#ROOT_DATE=281013
ROOT=time_lapse #assign root and the variable "num", e.g., 030314_$num
DIRS=([0]="X030914_BP2" [1]="X080914_BP1" [2]="X100914_BP1" [3]="X100914_BP2" [4]="X110914_BP1" [5]="X030914_P1" [6]="X030914_P2" [7]="X030914_P3" [8]="X080914_P2" [9]="X100914_P1_2" [10]="X030914_AP1" [11]="X030914_AP3" [12]="X100914_AP1" [13]="X110914_AP1" [14]="X030914_AP2_2") #this is where the fastq folders are
INPUT=/rdata/ngseq/original_data/rna/illumina/Guy/$ROOT #this is the parent directory of the folders "DIRS"
DATE=$(date +%d%m%y)
OUTPUT=/rdata/ngseq/Playground/guy/tophat2/$ROOT/$DATE
BT2INDEX=/rdata/ngseq/Playground/guy/bowtie2/Bowtie2Index_Arabidopsis/genome
OUT_LUC=/rdata/ngseq/Playground/guy/tophat2/$ROOT/$DATE"luc"
LUCINDEX=/rdata/ngseq/Playground/guy/bowtie2/luc_index/luc_index
th2path=/rdata/ngseq/Playground/guy/tophat2/$ROOT/$DATE
files=(${DIRS[@]/#/$th2path/})

for h in {0..14}; do
	FILES=("/rdata/ngseq/Playground/guy/Notes/At_ID_NAME" "${files[h]}""/""${DIRS[h]}"".txt")
done

#creating the directories . Note! if running twice at the same day, the first run will be overiden
for i in ${DIRS[*]}; do
	mkdir -p $OUTPUT/$i
done 

#running tophat
for l in ${DIRS[*]}; do
	tophat2 -N 1 -o $OUTPUT/$l -i 10 -p 2 --segment-mismatches 3 --min-segment-intron 10 --transcriptome-index=/rdata/ngseq/Playground/guy/tophat2/At_th2_index/At_th2_index $BT2INDEX $INPUT/$l/fastq/R1.fastq 2>&1 | tee -a $OUTPUT/$l/$l.log &
done
wait


#sorting by name with samtools
for m in ${DIRS[*]}; do
	samtools sort -n $OUTPUT/$m/accepted_hits.bam $OUTPUT/$m/accepted_hits_sort &
done
wait

#converting to sam files
for n in ${DIRS[*]}; do
	samtools view $OUTPUT/$n/accepted_hits_sort.bam > $OUTPUT/$n/accepted_hits_sort.sam &
done
wait

#counting w/ HTSeq
for p in ${DIRS[*]}; do
	python -m HTSeq.scripts.count -r name -s no -t exon -i gene_id -m intersection-strict $OUTPUT/$p/accepted_hits_sort.sam /rdata/ngseq/Playground/guy/Arabidopsis_thaliana_Ensembl/Ensembl/TAIR10/Annotation/Genes/genes_new.gtf > $OUTPUT/$p/$p"_1".txt &
done
wait

#Creating edgeR compatible text file (removing the last 5 summerizing records that HTSeq prints)
for q in ${DIRS[*]}; do
	awk 'NR<=33602' $OUTPUT/$q/$q"_1".txt > $OUTPUT/$q/$q.txt & # this command will print all first 33602 (all At#) and redirect it to a new file
done

##############################
#counting the luciferase reads
##############################
for i in ${DIRS[*]}; do
	mkdir -p $OUT_LUC/$i
done 

#running tophat
for l in ${DIRS[*]}; do
	tophat2 --no-novel-junc -o $OUT_LUC/$l $LUCINDEX $INPUT/$l/fastq/R1.fastq 2>&1 | tee -a $OUT_LUC/$l/$l.log &
done
wait

rm -r tmp.txt
for h in {0..14}; do
	FILES=(${files[h]}"/"${DIRS[h]}".txt")
	printf "%s\n" "${FILES[*]}" >> tmp.txt
done

FILES1=`less tmp.txt`
FILES2=(/rdata/ngseq/Playground/guy/Notes/At_ID_NAME ${FILES1[*]})
printf "%s %s %s\n" "At_num" "gene_name" "${DIRS[*]}" > /rdata/ngseq/Playground/guy/edgeR/summary_time_lapse.txt
paste ${FILES2[@]} | awk '{print $1,$2,$4,$6,$8,$10,$12,$14,$16,$18,$20,$22,$24,$26,$28,$30,$32}' >> /rdata/ngseq/Playground/guy/edgeR/summary_time_lapse.txt

#to understand this string manipulation see http://wiki.bash-hackers.org/syntax/pe#simple_usage
luc_files=(${DIRS[@]/#/$OUT_LUC/}) #this will substitute empty string before $DIRS with $OUT_LUC or in other words will create the paths $OUT_LUC/$DIRS
luc_files1=(${luc_files[@]/%//align_summary.txt}) #this will substitute the null string with at the end (%) of $luc_files a "/" (%/) and then align_summary.txt
DR5_EXP=([1]="DR5" [2]="DR5" [3]=`awk '/Mapped/ {print $2}' ${luc_files1[@]}`)
echo ${DR5_EXP[@]} >> /rdata/ngseq/Playground/guy/edgeR/summary_time_lapse.txt
sed 's/ /\t/g' /rdata/ngseq/Playground/guy/edgeR/summary_time_lapse.txt > /rdata/ngseq/Playground/guy/edgeR/counts_time_lapse.txt
