## supplemental figures and supplemental file description for "Cell wall remodeling and vesicle trafficking mediate the root clock in Arabidopsis"

Supplementary Materials

Figures S1-S7

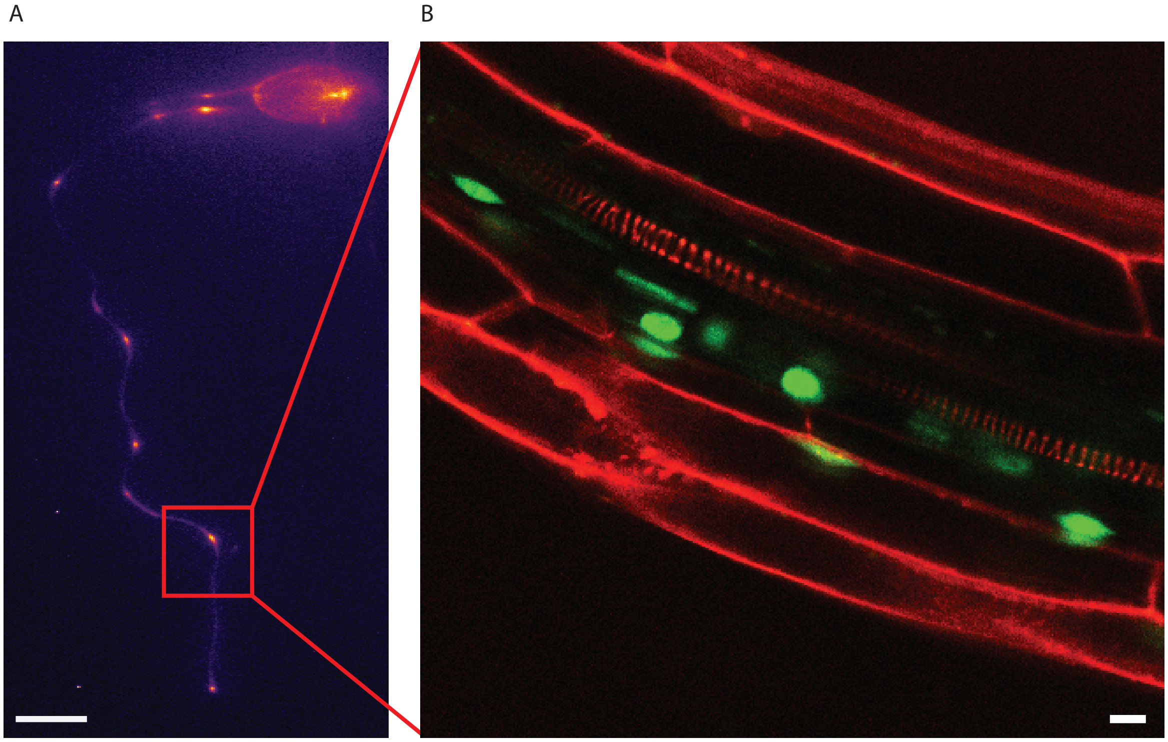


**Fig. S1. The youngest pre-branch site is associated with developing LRP. (A)** Lumazone imaging of DR5::LUC; pCLE44::GFP seedling. Youngest pre-branch site is marked by red square. Scalebar=0.2 cm. **(B)** same seedling as in (A). The region of the pre-branch site marked in (A) was bordered by proximal and distal lines and scored for pCLE44::GFP-marked LRP using the z-stack function of the confocal microscope. Scalebar=10 μm.

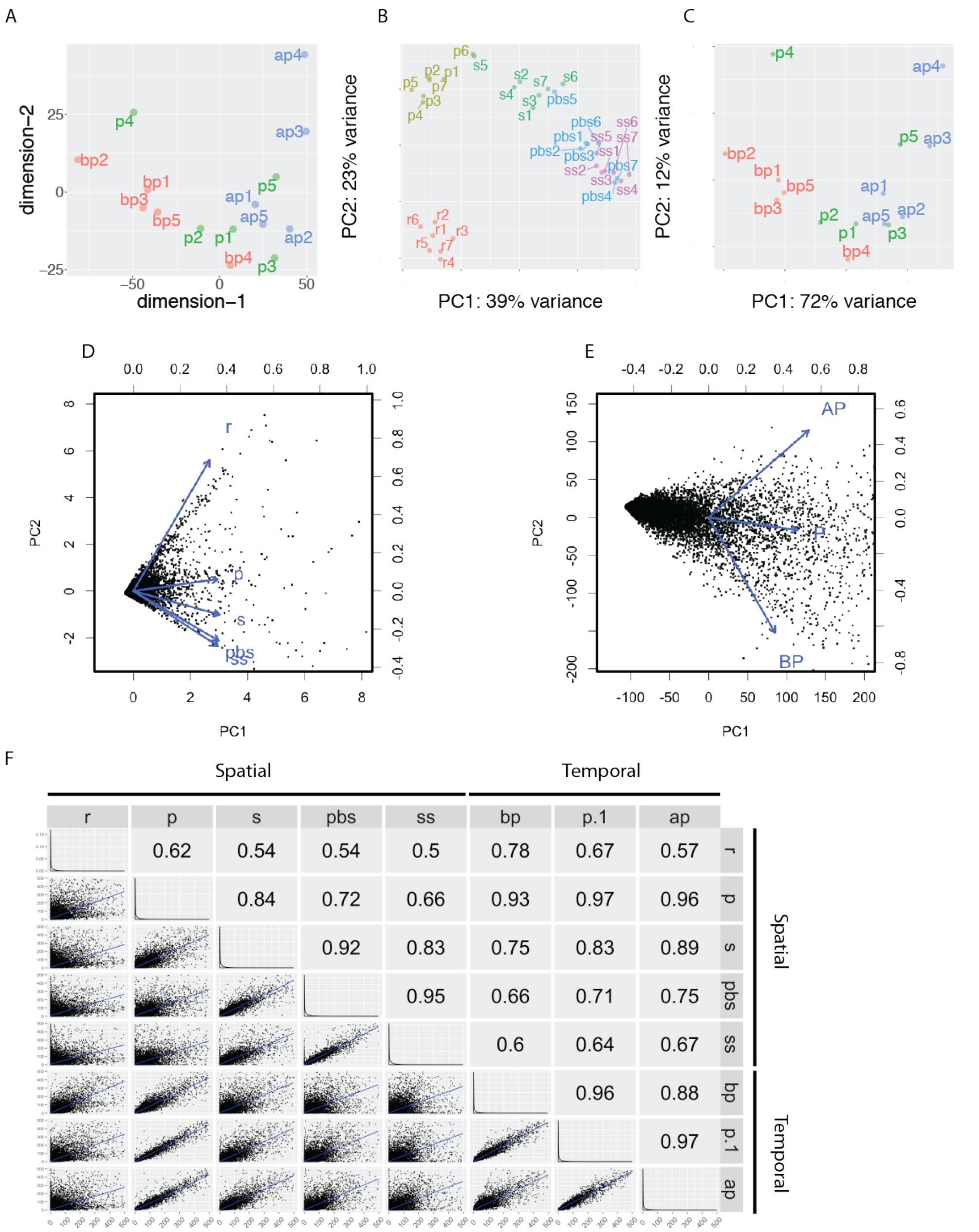


**Fig. S2. Statistical analysis of RNA-seq experiments confirms data quality. (A)** Multidimensional scaling analysis of the RNA-seq temporal experiment. **(B** and **C)** PCA analysis of the spatial (A) and temporal (B) experiments. Root section names correspond to those in Fig. 1, A and B, respectively. **(D** and **E)** Biplot analysis of the spatial (D) and temporal (E) experiments. Each point represents a single gene's coordinates projected onto the first two PCs. Each arrow represents a vector of the mean value of the indicated section. Note that the vectors are arranged (top to bottom, clockwise) in the same order as the sections along the root. Small angles between vectors e.g., the one between pbs and ss reflect high correlation between the regions while near-orthogonal vectors such as r and ss reflect uncorrelated relationships between samples. The biplots support the same trend shown in the MDS and PCA analyses. **(F)** Correlation analysis between all eight samples of the spatial and temporal experiments (p.1 is the p section of the temporal experiment). Correlation coefficients ($R^2$) are shown in the upper triangle of the plot. Note that p and p.1, which represent the OZ in the spatial and temporal experiments, respectively, have the highest $R^2$ value as expected.

A

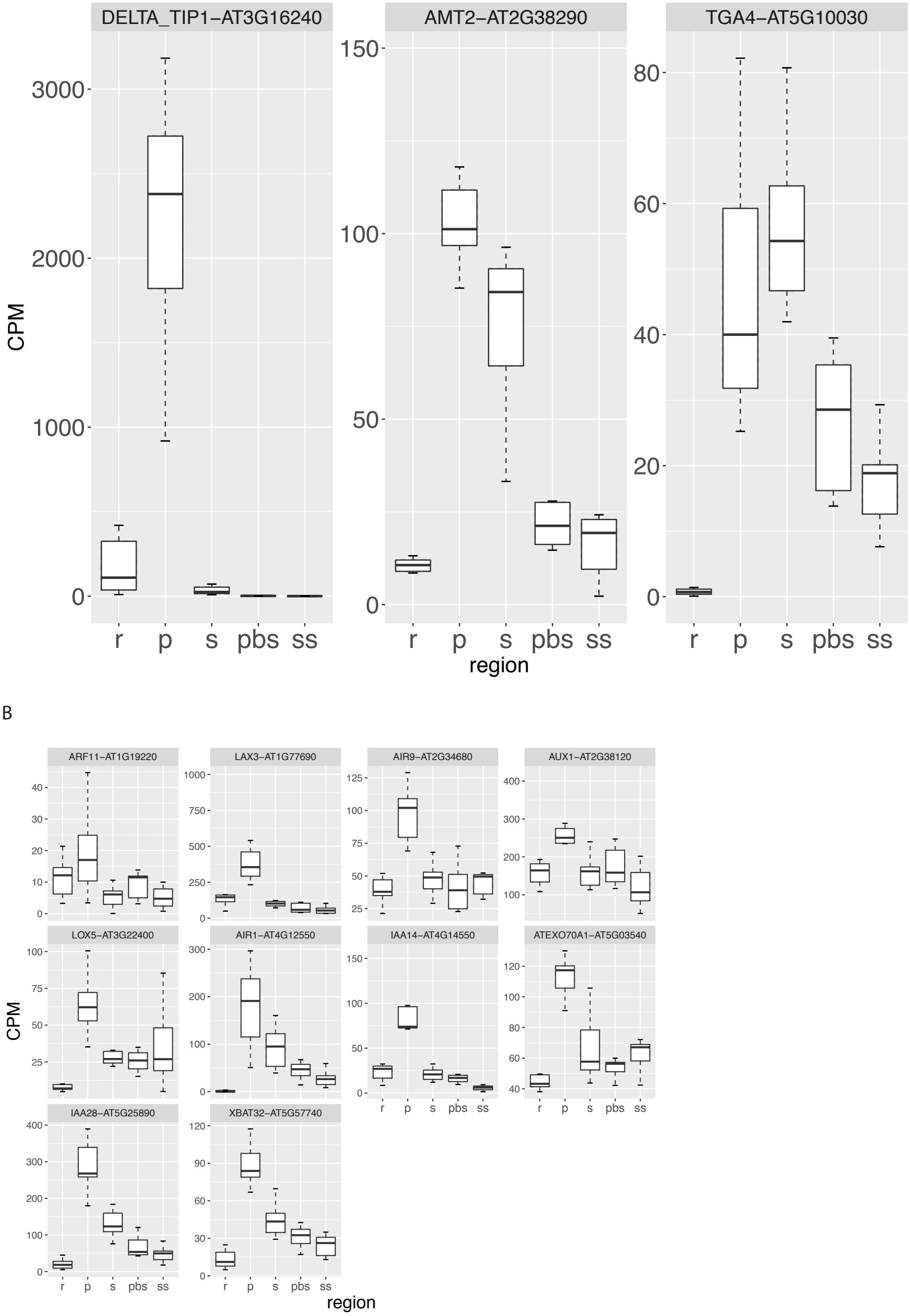


**Fig. S3. Expression of selected genes in the spatial experiment. (A)** Expression pattern of three nitrogen-related genes, which are also involved in LR initiation. **(B)** Expression pattern of the 10 genes in the LR GO term that had significant OZ expression in the spatial experiment.

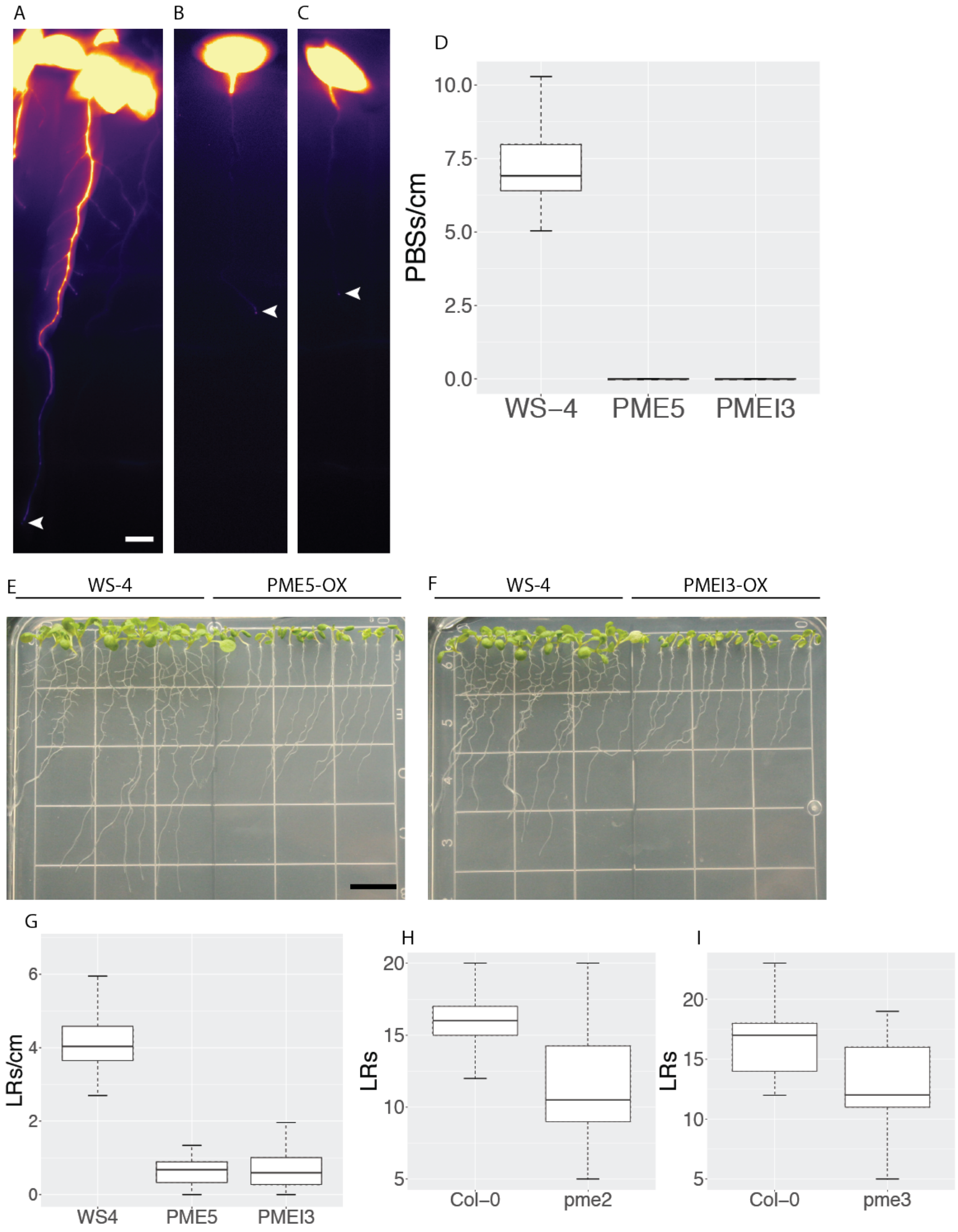


**Fig. S4. Destabilization of esterified and de-esterified pectin inhibits PBS and LR formation. (A** to **C)** WT DR5::LUC expressing seedlings (WS-4; (A)), PME5 overexpression (B) and PMEI3 overexpression (C) under EtOH inducible conditions. Arrowhead marks root tips. Scalebar=0.2 cm. **(D)** Quantification of DR5::LUC PBS number in WT (WS-4), PME5 overexpression and PMEI3 overexpression. p-value$<10^{-8}$ (Wilcoxon Rank Sum test). **(E** and **F)** Seedling phenotype of PME5 overexpression (E) and PMEI3 overexpression (F) compared to WS-4 WT. Scalebar=1 cm. **(E)** Quantification of LRs and LRP shown in (E and F). p-value$<10^{-10}$ (Wilcoxon Rank Sum test). **(H** and **I)** Quantification of LRs in *pme2* (F) and *pme3* (G). p-value=0.0002 (F) and p-value=0.01 (G) (Wilcoxon Rank Sum test).

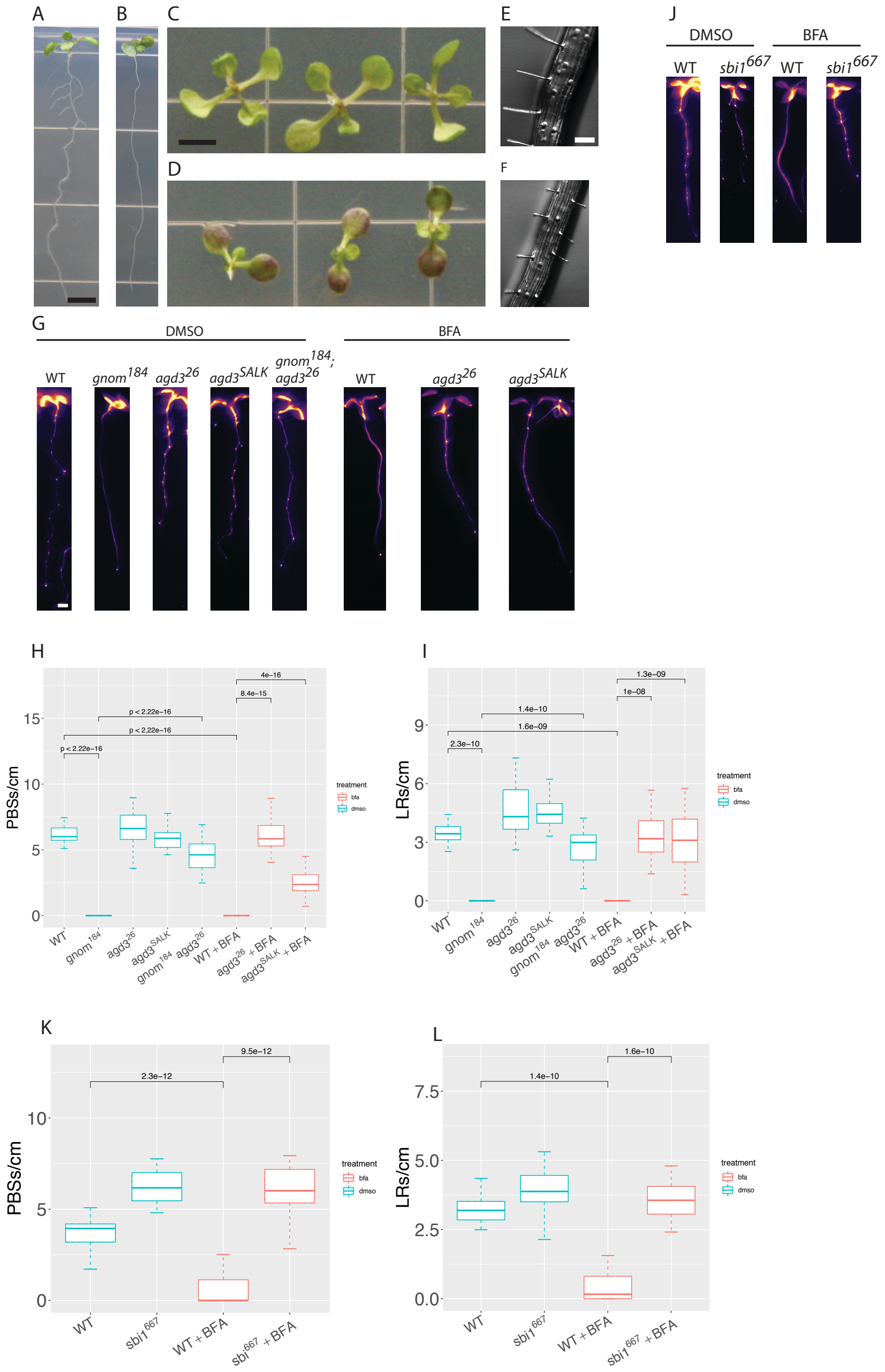


**Fig.S5. BFA treatment, *gnom*[184] and its suppressors affect lateral root development. (A and B)** WT (Col-0 (A)) and *gnom*[184] (B) seedling phenotypes. Scalebar=0.5 cm. **(C and D)** Shoot phenotype of WT (Col-0 (C)) and *gnom*[184] (D) seedlings. Scalebar=0.5 cm. **(E and F)** Root hair phenotype of WT (Col-0 (E)) and *gnom*[184] (F). Scalebar=100 µm. **(G)** DR5::LUC expression patterns. Scalebar=0.2 cm. **(H and I)** pre-branch site (H; DR5::LUC) and LR (I) count in the genotypes and treatments depicted in (G). p-values indicated above boxes. (t-test). **(J)** the given genotypes with and without BFA treatment. Scalebar=0.5 cm. **(K)** Quantification of pre-branch site (K) and LR (L) number in the *sbi*[667] mutant.

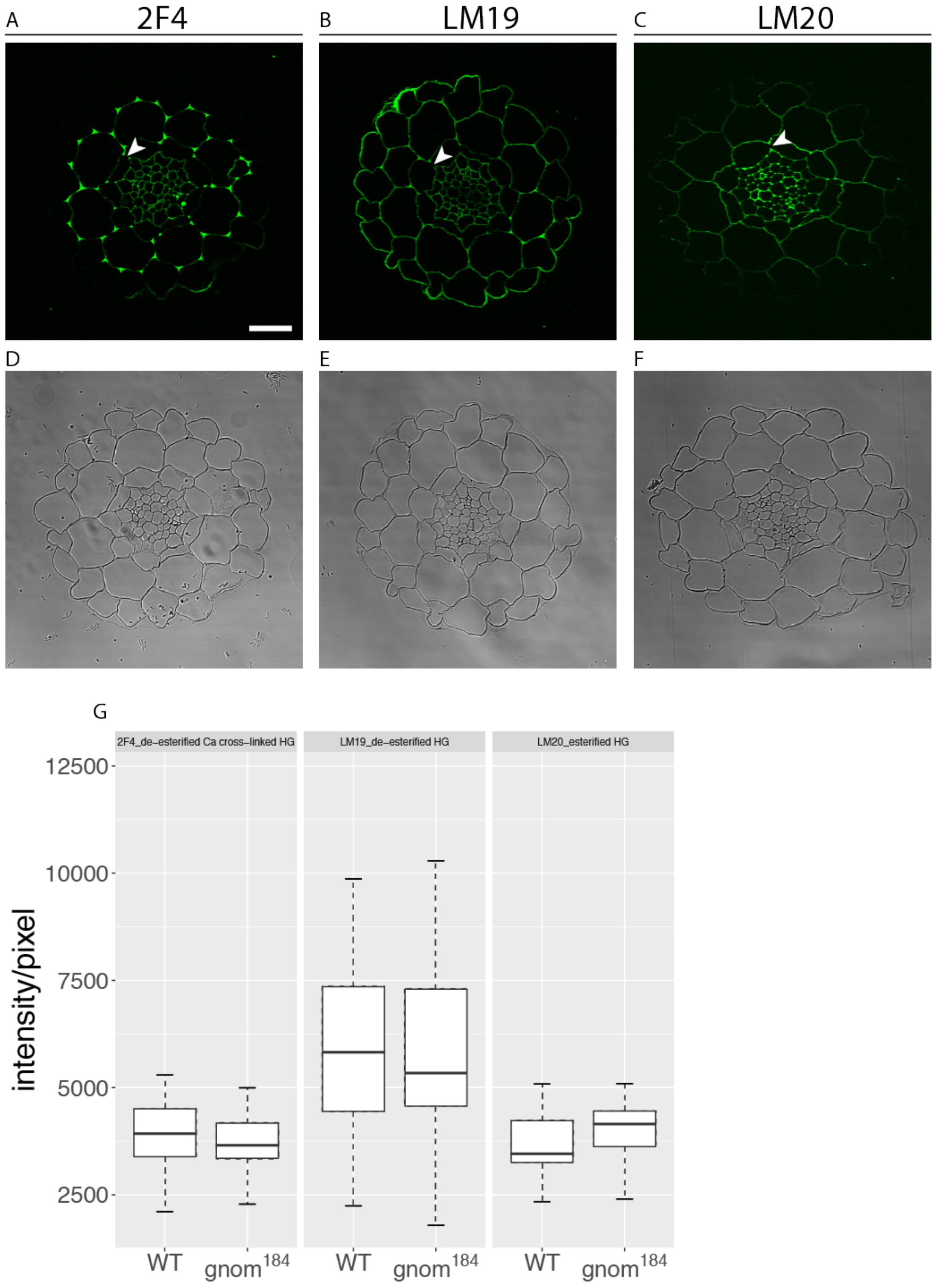


**Fig. S6. *gnom*$^{184}$ does not affect pectin density in the root. (A** to **C)** Immunohistochemistry of WT root cross sections labeled with 2F4 (A), LM19 (B) and LM20 (C) Abs. Note the lack of signal in Casparian strip position (arrowheads). **(D** to **F)** bright field images of the sections shown in (A to C). **(G)** Quantification of pixel intensity of the three Abs shown in (A to C) in WT and *gnom*$^{184}$ roots. Scalebar=20 μm.

A

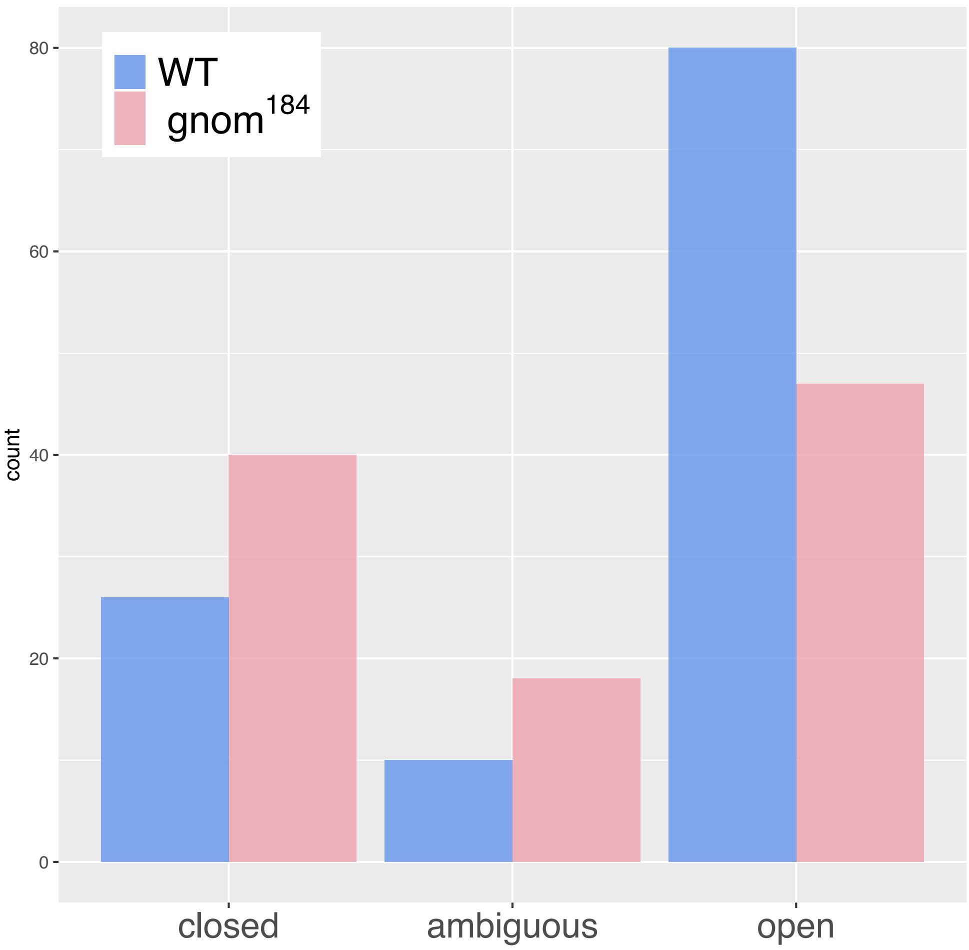


**Fig. S7. TEM qualitative valuation of de-esterified HG. (A)** Each pericycle-endodermis junction quantified in Fig. 4, A to C was evaluated as “open” where the gold particles form a near-triangle shape, “closed” with no clear space between the cell wall intersection or undetermined (ambiguous) in WT and *gnom*$^{184}$. The number of each type was plotted for each genotype. $p<0.002$ ($\chi^2$ test).

<u>Tables</u>

Table S1. RNA-seq read count results.
Table S2. Results of GO term analysis.
Table S3. EMS screens results.

Codes and text files

1. counts_spatial.txt. Read count of all 35 spatial experiment libraries.
2. counts_temporal.txt. Read count of all 15 temporal experiment libraries.
3. TAIR10_functional_descriptions.txt. TAIR10 gene list with functional description.
4. cell_wall_inten_ijf.txt. Fiji macro for extracting the area and intensities of Abs marked regions (related to Fig. S6).
5. fiji-count_gold_particles_per-area.txt. Fiji macro for calculating gold particles per area (related to Fig. 4, B to D).
6. threshold_measure_wall_intensity_raul.txt. Fiji macro for selecting and measuring cell wall areas with Ab fluorescence (related to Fig, 4A).
7. Ab_lrp.R. R code used to calculate p-values and draw plots of Ab intensity in regions of LRP emergence (related to Fig. 4, G and H).
8. TEM_measurement.txt. R code used to measure pecti-marked gold particles per area (related to Fig. 4D).
9. counts_spatial.txt. Count number of the spatial experiment. Each of the seven roots (digits 1-6 and the last one) and the five section types (R, P, S, PBS and SS) is given in the header.
10. counts_temporal.txt. Count number of the temporal experiment.
11. luc_count_spatial.sh. Bash code for counting Luciferase reads in the spatial experiment.
12. pipeline_spatial.sh. Bash script that takes fastq files of the spatial RNA-seq experiment and output the read count (given in the counts_spatial.txt file).
13. pipeline_temporal (including luc reads and binding).sh. Same as previous for the temporal experiment but also counts and merges the Luciferase reads.
14. TAIR10_functional_descriptions.txt.
15. analysis_spatial.R. R code for analyzing and ploting the RNA-seq spatial experiment.
16. analysis_temporal.R. Same as previous for the temporal experiment (need to be ran before analysis_spatial.R).
17. gprofiler.R. R code for analyzing and ploting GO term analysis.
